# Molecular dynamics and theratyping in airway and gut organoids reveal R352Q-CFTR conductance defect

**DOI:** 10.1101/2021.08.11.456003

**Authors:** Sharon L. Wong, Nikhil T. Awatade, Miro A. Astore, Katelin M. Allan, Michael J. Carnell, Iveta Slapetova, Po-chia Chen, Jeffry Setiadi, Elvis Pandzic, Laura K. Fawcett, John R. Widger, Renee M. Whan, Renate Griffith, Chee Y. Ooi, Serdar Kuyucak, Adam Jaffe, Shafagh A. Waters

**Author notes:** These authors contributed equally to this work (Sharon L. Wong, Nikhil T. Awatade, Miro A. Astore). School of Natural Sciences (Chemistry), University of Tasmania (Renate Griffith). **Authors’ contributions** Conception and design: SAW, AJ. Recruiting of participants: LF, SAW, JW, CYO. Collection of nasal brushings: LF, JW. hNECs culture and ion transport assay and analysis: NTA, KA, SLW. Immunofluorescent microscopy: SLW. CBF imaging: IS. CBF analysis design: EP. Collection of rectal biopsies: CYO. Culturing the organoids: NTA, SLW, SAW. Performing FIS: IS, NTA, KA. FIS assay scripts: MC. FIS assay analysis: NTA, MC. Molecular Dynamics: MA, PC, JS, RG, SK. Figure preparation: SLW, KA, MA, SAW. Writing – original draft: SLW, MA, SAW. Review and editing: SAW, KA, LF, RG, SK with intellectual input from all other authors. Supervision: SAW, SK. **Correspondence:** Dr. Shafagh Waters.

## Abstract

A significant challenge to making targeted CFTR modulator therapies accessible to all individuals with cystic fibrosis (CF) are many mutations in the CFTR gene that can cause CF, most of which remain uncharacterized. Here, we characterized the structural and functional defects of the rare *CFTR* mutation R352Q – with potential role contributing to intrapore chloride ion permeation – in patient-derived cell models of the airway and gut. CFTR function in differentiated nasal epithelial cultures and matched intestinal organoids was assessed using ion transport assay and forskolin-induced swelling (FIS) assay respectively. Two CFTR potentiators (VX-770, GLPG1837) and a corrector (VX-809) were tested. Data from R352Q-CFTR were compared to that of participants with mutations with known impact on CFTR function. R352Q-CFTR has residual CFTR function which was restored to functional CFTR activity by CFTR potentiators but not the corrector. Molecular dynamics (MD) simulations of R352Q-CFTR were carried out which indicated the presence of a chloride conductance defect, with little evidence supporting a gating defect. The combination approach of *in vitro* patient-derived cell models and *in silico* MD simulations to characterize rare *CFTR* mutations can improve the specificity and sensitivity of modulator response predictions and aid in their translational use for CF precision medicine.

## Introduction

Cystic fibrosis (CF), a rare life-limiting disorder affecting ~90,000 individuals worldwide [1], results from malfunction in the CF transmembrane conductance regulator (CFTR) protein. CFTR functions as a cAMP stimulated anion channel [2]. Chloride ions diffuse through the CFTR channel pore, causing a charge imbalance that is corrected by the flux of sodium ions through sodium channels [2]. The resultant ion imbalance causes osmosis of water into the extracellular space. Over 2000 variants in the *CFTR* gene are known. *CFTR* mutations are grouped into six major classes based on their functional defects in protein synthesis (I), intracellular maturation, processing or folding (II), channel gating (III) or conductance activity (IV), diminished quantity (V) and diminished stability (VI) [3].

CFTR modulators can restore mutant CFTR protein function [4]. Two subclasses of modulators are in current clinical use, though approval varies between countries. Potentiators such as ivacaftor (VX-770|Kalydeco®) increase CFTR channel gate opening. Correctors such as lumacaftor (VX-809), tezacaftor (VX-661) and elexacaftor (VX-445), increase delivery of misfolded CFTR to the cell surface. Combination therapies of potentiator and corrector(s) (Orkambi®, Symdeko®, Trikafta®) are approved for use in CF patients homozygous for DF508, the most common *CFTR* mutation [5–7]. The remaining CF patients are either compound heterozygous or homozygous for mutations other than DF508, only some of which are approved for treatment with modulator therapy [8]. A large number of *CFTR* mutations are not approved for modulator therapy because their exact mechanism of CFTR dysfunction and/or responsiveness to modulator therapy is often unknown. Traditional randomized clinical trials of modulators to identify modulator-responsive CF patients with rare *CFTR* mutations are costly, time-consuming and impractical [9].

A major breakthrough in the field of CF has been the development of two pre-clinical patient- and organ-specific cell models [10–12]. Both models are used in functional assays that allow rapid quantitative measurements of CFTR function. Differentiated airway epithelial cells are used in transepithelial ion transport assays (Isc) [13]. Intestinal organoids are used in forskolin (Fsk)-induced swelling (FIS) assays [14]. Each serves as a personalized functional model of the patient’s *CFTR* mutation and response to modulators. However, how cell models of the airway compare to those of the gut created from the same patient is not well understood. Mean changes in CFTR function *in vivo* correlate with the CFTR rescue assessed via the Isc and FIS assays in cell models of subjects with the same mutations [11, 13]. In addition, cell model responses in subjects with ultra-rare mutations have successfully predicted clinical benefit [15–17]. Results so far support the use of patient-derived cell models to guide personalized treatment [18].

We established human nasal epithelial cultures (hNECs) from ten participants with wild-type functioning CFTR (WT/WT) and seven participants with CF. The CF participants included five participants with known and characterized CFTR folding/maturation defects (DF508/DF508) and one with a CFTR gating defect (G551D/DF508). The result from these *CFTR* mutations with a known impact on CFTR function were used as reference and were compared to the result from one participant with the R352Q/DF508 *CFTR* genotype. We also created matched intestinal organoids from all seven participants with CF (**Supplementary material E1**). R352Q, a rare *CFTR* mutation in the CFTR channel pore, was chosen as it has been characterized in heterologous systems but is yet to be tested in a patient-derived model. Single channel and whole cell patch-clamp studies have demonstrated the R352 to be a molecular determinant of anion selectivity and permeability in the CFTR channel pore [33]. Our aims were to assess CFTR baseline activity and CFTR modulator response in I_sc_ and FIS assays. Potentiator agents VX-770 and GLPG1837 (a potentiator in clinical trials; hereafter G1837) were tested individually and in combination with a CFTR protein-folding corrector VX-809. In addition, to gain complementary and a better understanding of the effect of R352Q mutation on CFTR function, *in silico* structural atomistic modelling and molecular dynamics simulations were performed.

## Materials and Methods

Materials and methods are available in **Supplementary material E2**.

## Results

### R352Q-CFTR baseline activity and response to CFTR modulators in nasal epithelial cells

Mature, differentiated hNECs were pseudo-stratified (**Fig 1A**) and had functional beating cilia (CBF=6.2±0.1 Hz; **Fig 1B, Supplementary material E4**) and intact junction integrity (**Fig 1A**, TEER≥170 Ω.cm^2^; **Supplementary material E3A**). To assess ion transport, short-circuit current (Isc) measurements were performed (**Supplementary material E3B-D**). We first determined CFTR activity threshold in reference cell models which were used for comparison to R352Q-CFTR functional activity before and after modulator stimulation. Forskolin-stimulated CFTR-dependent anion currents (ΔI_sc-fsk_) in WT/WT hNECs were 21.2±1.2 μA/cm^2^ at baseline (**Fig 1C-D, Supplementary material E5**). Baseline ΔI_sc-fsk_ in DF508/DF508 hNECs were at 3.4±0.5 μA/cm^2^ and were not increased beyond ~0.8 μA/cm^2^ with either potentiator (**Fig 1D)**. We considered there to be an equal protein expression level from each of the DF508 alleles and therefore attributed each allele to equally contribute 0.4 μA/cm^2^ to potentiator-stimulated currents. This value is used to estimate the contribution of the DF508 allele to the experimental I_sc_ data of the heterozygous R352Q/DF508 participant. The reference gating G551D/DF508 hNECs had ΔI_sc-fsk_ of 4.2±0.8 μA/cm^2^ at baseline. Treatment with either potentiator caused at least a 2-fold increase in CFTR activity above baseline (net increase: VX-770: 4.2 μA/cm^2^; G1837: 8.2 μA/cm^2^) (**Fig 1D**).

**Figure 1.**
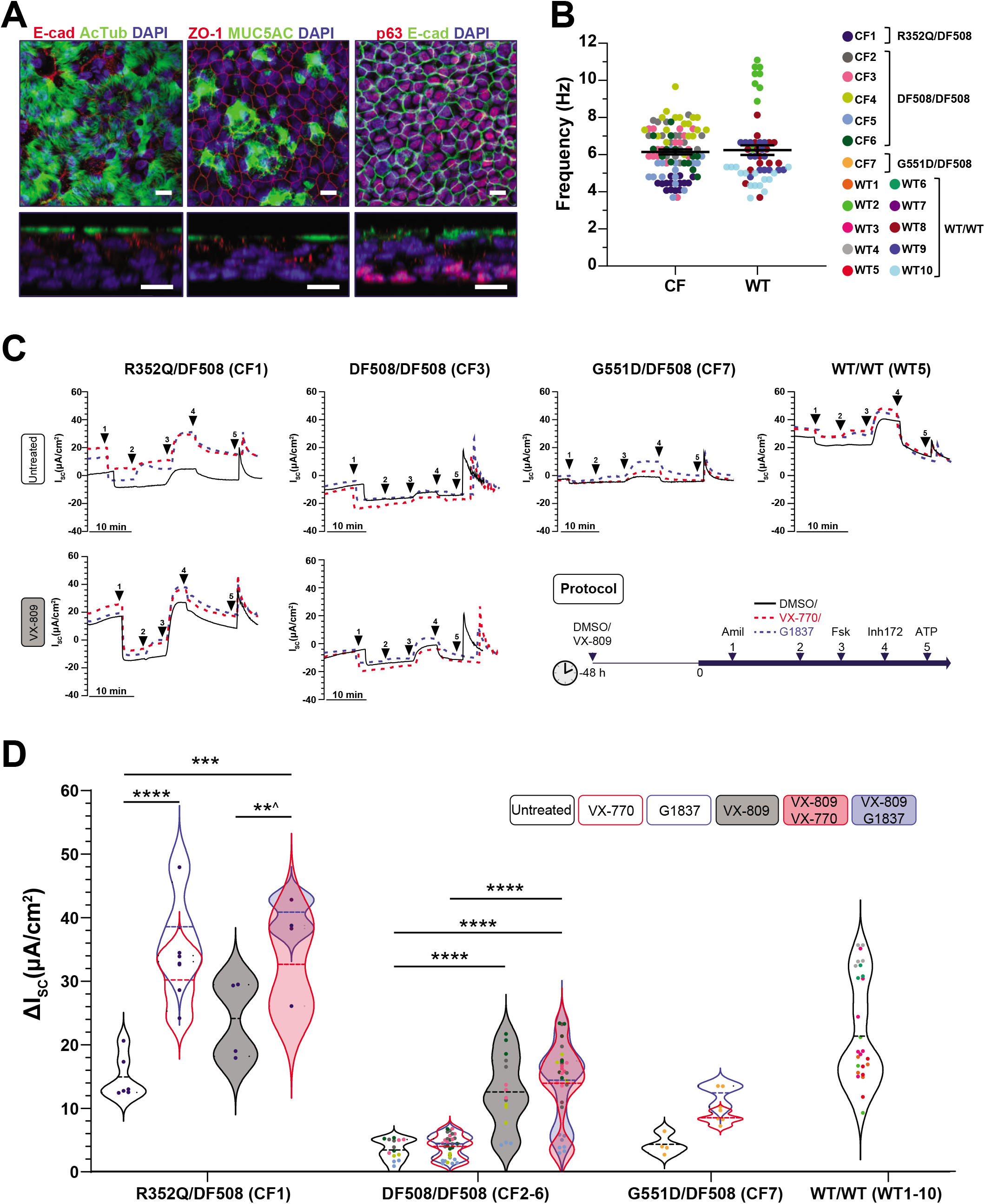
Functional response of R352Q-CFTR to CFTR modulators in differentiated human Nasal Epithelial Cells (hNECs). (A) Immunofluorescence staining of acetylated tubulin (ciliated cells), MUC5AC (goblet cells) and p63 (basal progenitor cells) in hNECs derived from a R352Q/DF508 participant (CF1). E-cadherin (adherens junction) and ZO-1 (tight junction) are localised to the intercellular junctions of epithelial cells. The top panels are top-view while the bottom panels are side-view showing the pseudostratified epithelium. 63x/1.4 oil immersion objective. Scale bars = 10 μm. (B) Dot plots of mean Cilia Beat Frequency (CBF) measurements (Hz) of the fully differentiated mature hNECs from CF and non-CF (WT) participants. CFTR genotypes of individuals are indicated. Each participant is coded with a different colour (legend in the figure). Each dot represents a single field of view of CBF measurement. (C) Representative Ussing chamber recordings of short circuit current (Isc) in hNECs from participants with CF and WT-CFTR control participants. Protocol used to measure functional CFTR expression in hNECs in 0.01% DMSO vehicle (untreated; top) or pre-treated with corrector (3 μM VX-809 for 48 h; bottom) followed by sequential addition of 100 μM apical amiloride (1. Amil), apical addition of either vehicle control 0.01% DMSO (black), or 10 μM VX-770 (red) or 10 μM G1837 (blue) (2. DMSO, VX-770, G1837), 10 μM basal forskolin (3. Fsk), 30 μM apical CFTR inhibitor (4. CFTRinh-172) and 100 μM apical ATP (5. ATP). A basolateral-to-apical chloride gradient was used. (D) Violin plots of total currents stimulated by DMSO or VX-770 or G1837 plus forskolin in hNECs untreated or pre-treated with VX-809. Data are from one R352Q/DF508, five DF508/DF508, one G551D/DF508 participants with CF and ten WT-CFTR control participants. n, number of participants. Each participant is coded with a different colour. Each dot represents an individual replicate. Data represented as violin plots to show distribution of data. One-way analysis of variance (ANOVA) was used to determine statistical differences. ***P < 0.001 and ****P < 0.0001. **^P < 0.01 for G1837 only. Where VX-770 and G1837 (overlapping violin plots) both achieved statistical significance, the least significant P-value is shown.

R352Q/DF508 hNECs demonstrated baseline CFTR activity of 14.8±1.4 μA/cm^2^, an appreciable 70% of WT-CFTR activity (**Fig 1C-D**). Potentiation with VX-770 or G1837 led to a significant (P<0.0001) 2-fold increase in CFTR activity, reaching 15.2 and 23.5 μA/cm^2^ above baseline respectively (~140–180% WT-CFTR activity). This response is similar to the 2-fold potentiator-stimulated response observed in the reference gating G551D/DF508 hNECs. The majority of the response to potentiators is most likely contributed by the restored R352Q because the total response is far greater than the 0.4 μA/cm^2^ which we have attributed to that of the DF508 response (**Fig 1D**). We conclude that R352Q-CFTR has high baseline activity amenable to potentiator rescue in hNECs.

We next explored whether corrector monotherapy or co-therapy with potentiators increased R352Q-CFTR functional rescue. In DF508/DF508-CFTR cultures pre-treated with VX-809, fsk alone significantly (P<0.0001) enhanced CFTR-mediated Cl^−^ currents by 3.7-fold, reaching 9.0 μA/cm^2^ above baseline (**Fig 1C-D**). We attributed each of the DF508 alleles to equally contribute 4.5 μA/cm^2^ to this increase in current. VX-809 co-therapy with either VX-770 or G1837 significantly (P<0.0001) increased the currents stimulated by potentiator monotherapy by approximately 10 μA/cm^2^ (**Fig 1D**). Unlike DF508/DF508-CFTR, the 9.1 μA/cm^2^ increase in R352Q/DF508 CFTR activity from the baseline in response to VX-809 monotherapy was not statistically significant (P=0.09). VX-809 co-therapy with either potentiator also resulted in a non-significant increase in VX-770- or G1837-potentiated CFTR currents by ~2 μA/cm^2^ (**Fig 1D**). Since VX-809 does not rescue R352Q, then R352Q-CFTR is unlikely to have a folding/maturation defect. The modest increase in current with VX-809 treatment was mostly, if not fully imparted by the rescued DF508 allele in the R352Q/DF508 hNECs.

### R352Q-CFTR baseline activity and response to CFTR modulators in intestinal organoids

We evaluated CFTR activity in matched intestinal organoids from the same participant with the R352Q/DF508 genotype and those participants with reference DF508/DF508 and G551D/DF508 genotypes using a fsk-induced swelling (FIS) assay. To detect baseline and modulator-stimulated CFTR activity in the organoids, swelling was assessed at four fsk concentrations ranging from 0.02 to 5 μM. FIS of organoids was dependent on fsk dose, *CFTR* genotype and the individual participant (**Fig 2A**). At 0.8 μM fsk – the optimal concentration for baseline FIS assessment [11] – minimal swelling was observed for the DF508/DF508 (AUC=42.8±19.4) and G551D/DF508 organoids (AUC=82.9±20.1) (**Fig 2A**). R352Q/DF508 organoids showed considerably higher FIS at the same 0.8 μM fsk concentration (AUC=196.3±19.9) (**Fig 2A**).

**Figure 2.**
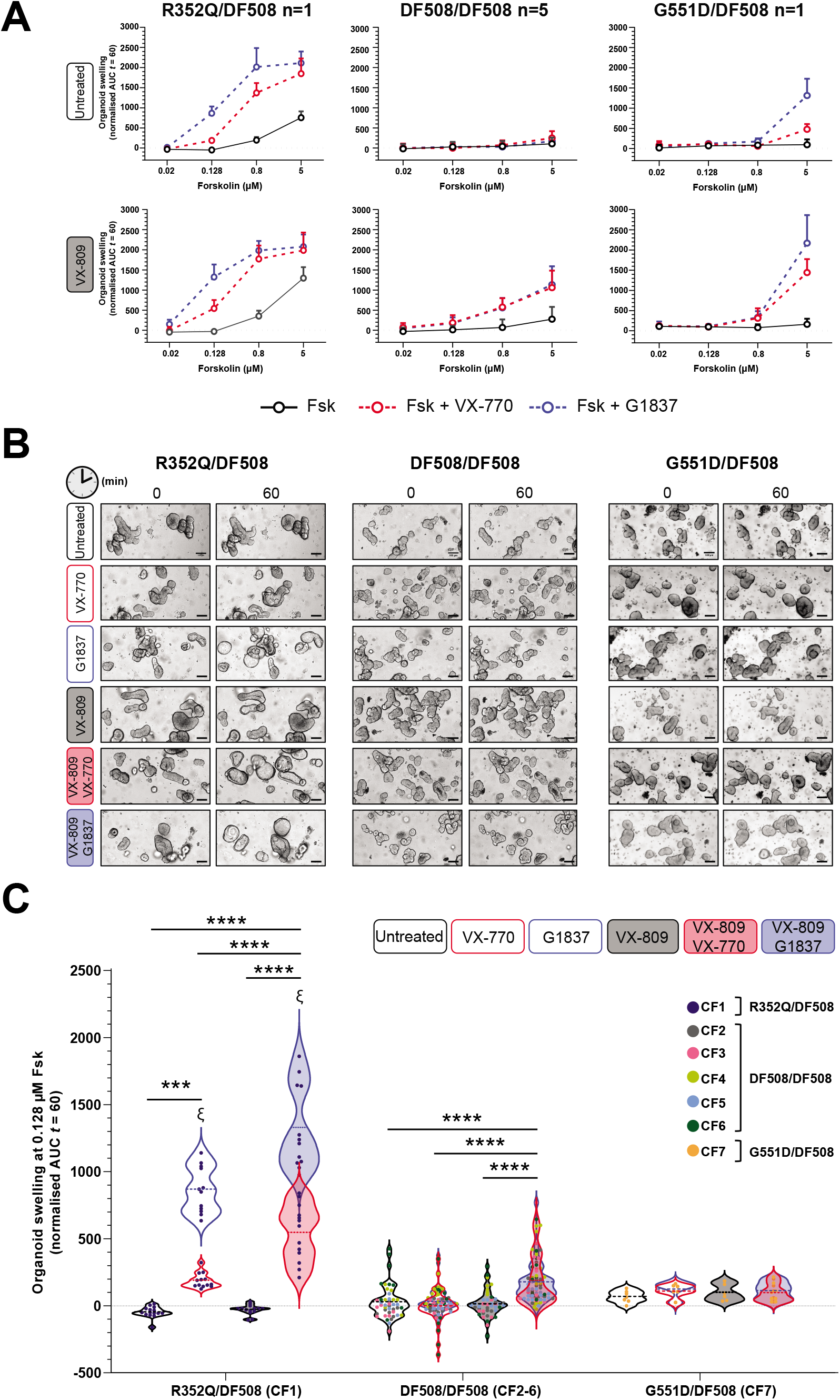
Functional response of R352Q-CFTR to CFTR modulators in intestinal organoids. Forskolin-induced swelling (FIS) assay in organoids (left to right) from one R352Q/DF508, five DF508/DF508 and one G551D/DF508 participants with CF. Organoids were incubated overnight with 0.03% DMSO (vehicle) or 3 μM VX-809. After 24h, a range of forskolin concentrations from 0.02 to 5 μM, were acutely added, either alone or in combination with 3 μM VX-770 (red) or 3 μM G1837 (blue). (A) Values plotted are the means ±standard deviation (SD) of the area under the curve (AUC). (B) Representative brightfield images of organoids from each CFTR genotype at baseline (t=0) and after 1h of stimulation (t=60) at 0.128 μM forskolin. Scale bars = 100 μm. (C) Violin plots of FIS response (AUC at 0.128 μM forskolin) of organoids from participants with CF, expressed as the absolute AUC calculated from the time periods t=0 (baseline) to t=60. Data represented as violin plots to show distribution of data. Each participant is coded with a different colour (legend in the figure). Each dot represents an individual replicate. n, number of participants. One-way analysis of variance (ANOVA) was used to determine statistical differences. ***P < 0.001 and ****P < 0.0001. ξP < 0.0001 for VX-770 vs G1837. Where VX-770 and G1837 (overlapping violin plots) both achieved statistical significance, the least significant P-value is shown.

FIS of intestinal organoids at 0.128 μM fsk has been demonstrated to correlate strongly with clinical modulator responses and therefore is used to assess CFTR modulator response *in vitro* [11]. At this fsk concentration, organoids – independent of *CFTR* genotype – did not demonstrate FIS greater than AUC of 67.1 (**Fig 2B-C, Supplementary material E6**). In DF508/DF508 organoids, treatment with either VX-770 or G1837 did not induce any swelling, suggesting no improvement in CFTR activity with potentiator therapy. G551D/DF508 organoids showed a modest 1.5-fold increase in FIS following treatment with either potentiator, increasing AUC by 38.9 (VX-770) and 54.2 (G1837) above baseline (**Fig 2B-C)**. In R352Q/DF508 organoids, potentiation with either VX-770 or G1837 led to a significant (P<0.001) increase in FIS, increasing AUC by 234.6 and 910.6 above baseline respectively (**Fig 2B-C**). G1837 was significantly (P<0.0001) more effective in increasing FIS when compared to VX-770 in R352Q/DF508 organoids, but not in organoids with the other *CFTR* genotypes wherein the efficacy of the potentiators did not significantly differ from each other (**Fig 2C, Supplementary material E6**). These data indicate that R352Q has residual CFTR function and is responsive to VX-770 and G1837.

We next determined the effect of VX-809 monotherapy and co-therapy in R352Q/DF508 organoids. In reference DF508/DF508 and G551D/DF508 organoids, VX-809 monotherapy at 0.128 μM fsk concentration did not yield a significant change in FIS, with net increase in AUC of ≤30.4 above baseline (**Fig 2B-C, Supplementary material E6**). In DF508/DF508 organoids, VX-809 co-therapy with either VX-770 or G1837 caused a significant (P<0.0001) increase in FIS compared to potentiator monotherapy, with net increase in AUC of 189.3 and 145.6 respectively (**Fig 2B-C**). In contrast, no additional increase in FIS was observed in G551D/DF508 organoids following co-therapy with VX-809 and either potentiator (**Fig 2B-C**). Similar to the reference DF508/DF508 organoids, no change in FIS was observed in the R352Q/DF508 organoids at 0.128 μM fsk with VX-809 monotherapy. The net increase in AUC was 17.7 above baseline (**Fig 2B-C**). VX-809 co-therapy in R352Q/DF508 organoids followed the same trend as the DF508/DF508 organoids but with a larger magnitude of response. VX-809 co-therapy with VX-770 or G1837 significantly (P<0.0001) increased FIS beyond each respective monotherapy by 355.2 and 459.7 (**Fig 2B-C**).

### Correlation of CFTR modulator-response in hNECs with participant-matched intestinal organoid swelling assay outcomes

We compared the CFTR modulator response as determined by ion transport in hNECs and FIS in intestinal organoids. To determine the amount of modulator-stimulated restoration above the baseline CFTR activity, we subtracted the ΔI_sc_/AUC values with fsk alone from those with fsk plus modulators. A linear positive correlation was observed (r=0.75; P<0.0001; **Fig 3A**). Participant-to-participant variability was evident (**Fig 3B, Supplementary material E7**). Amongst the modulators, an inverse correlation was observed in response to VX-809 monotherapy between the two cell models (r=−0.74, **Fig 3C**). Whereas VX-809 monotherapy significantly increased baseline currents in hNECs, no increase in FIS was detected in the matched organoids (**Supplementary material E7**).

**Figure 3.**
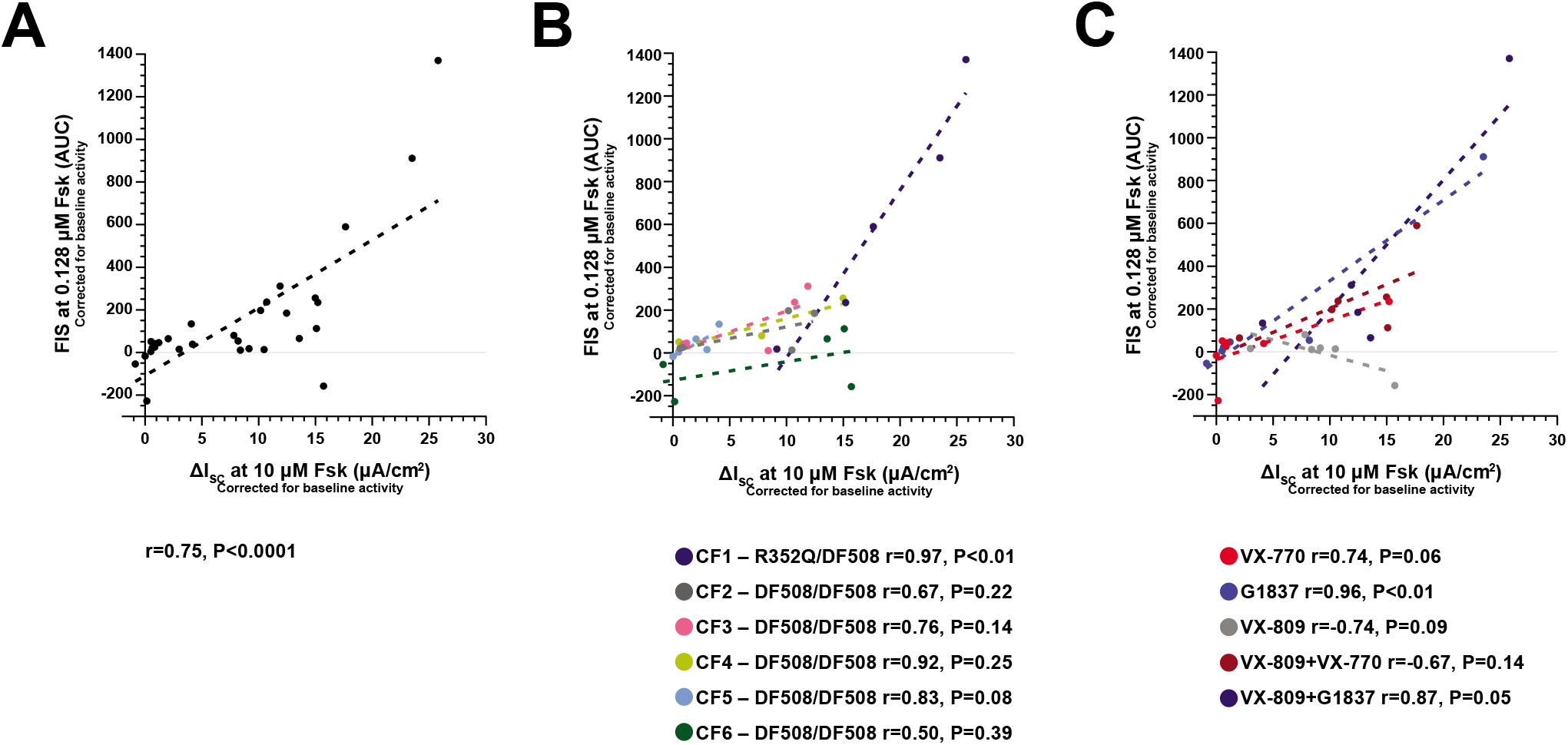
Correlation between individual in vitro treatment-modulated CFTR function in participant-matched differentiated human Nasal Epithelial Cells (hNECs) and intestinal organoids. Modulator-enhanced CFTR response – corrected for the baseline CFTR activity – in hNECs (short circuit current, ΔIsc) compared with organoids (forskolin-induced swelling, FIS) from each participant shown (A) as a collective, (B) by participant and (C) by modulator therapy. The dotted line through data points shows simple linear regression analysis. Correlation (r) values were calculated using Pearson linear correlation.

### MD simulations and free energy calculations reveal the dysfunction of the R352Q mutant

In line with the previously proposed role for R352 in forming a salt bridge [19], it was observed that mutation of the positively charged arginine (R) to neutral glutamine (Q) disrupted the R352-D993 salt bridge in the midsection of the CFTR channel pore (**Fig 4A, Supplementary material E8**). This salt bridge is a key structural element in CFTR, and its disruption has been shown to affect properties of both conductance and gating [19]. On the other hand, in the microsecond long MD simulations, the root-mean-square deviations (RMSDs) of the R352Q-CFTR membrane spanning helices were stable, indicating no discernible disruption to the channel pore architecture (**Fig 4B**). The data suggest that the R352Q-CFTR open conformation can remain stable on the sub-microsecond time scale necessary for chloride conduction [20]. If gating was impacted by the R352Q mutation, longer simulations might be able to decipher any disrupted pore architecture.

**Figure 4.**
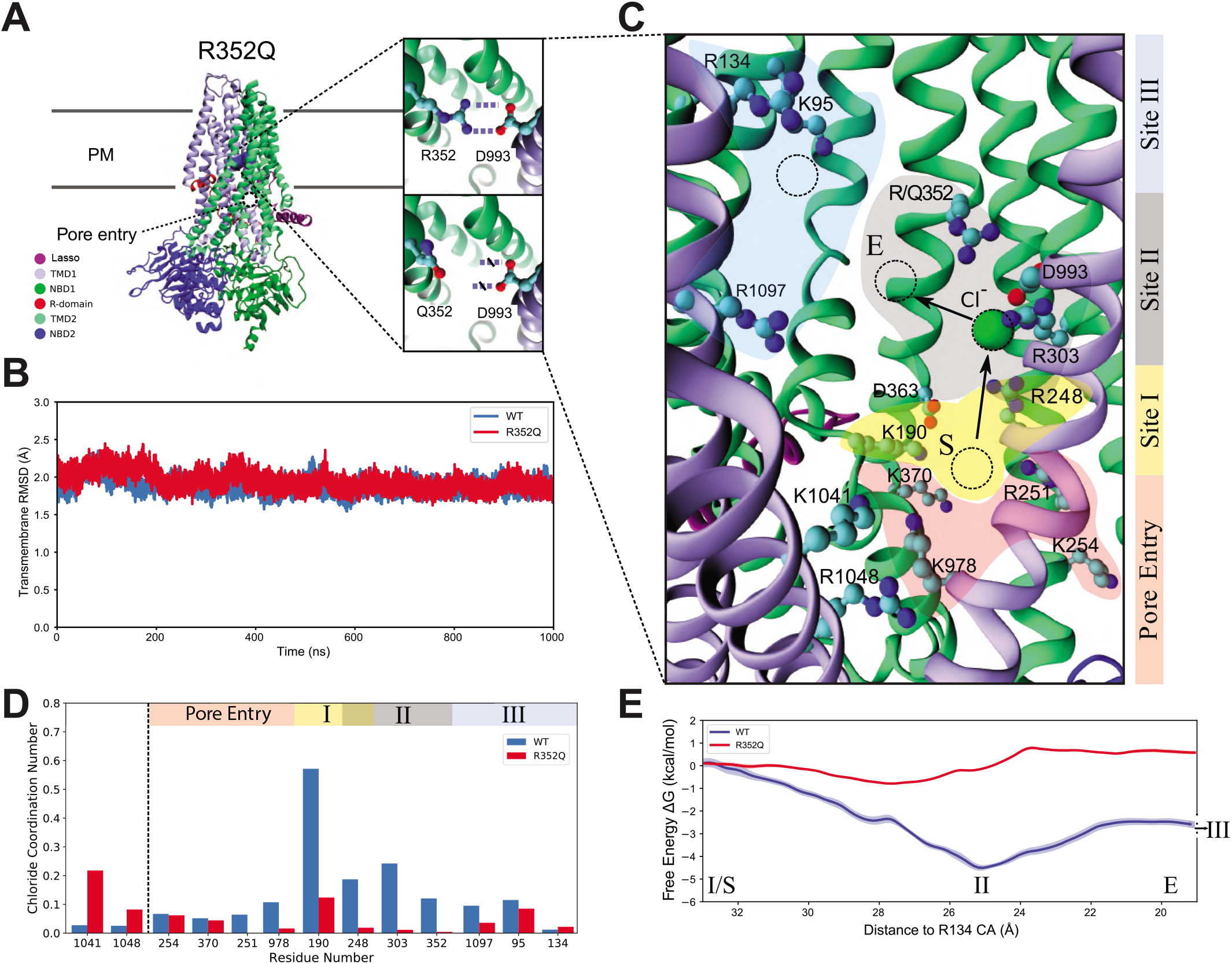
Simulations provide a mechanistic understanding of R352Q-CFTR dysfunction. (A) Atomistic structure of the CFTR channel pore based on cryo-EM (6MSM), with the configuration of the R352-D993 salt bridge and the disrupted R352Q highlighted in insets. Mutation of the charged R352 side chain (right-top) to the neutral Q352 side chain (right-bottom) disrupts the R352-D993 salt bridge present in WT-CFTR. (B) The root-mean-square deviations (RMSDs) of the transmembrane domains of WT (blue) and R352Q (red) -CFTR channel pore, computed using the 6MSM coordinates as reference. (C) A visualisation of charged residues that line the CFTR channel pore. The path of chloride ion conduction from pore entry (pink) to site I (yellow), sites II (grey) and III (blue) is indicated. The computed free energy path in (E) is defined with reference to the chloride ions distance to the alpha carbon of R134 in site III, with the start and end points delineated by S and E respectively. Charged residues lining the channel pore are shown as ball and sticks. An example chloride ion is represented with a green ball in site II. (D) The histogram showing the chloride ion occupancy in WT (blue) and R352Q (red) -CFTR. The number of chloride ions within 4 Å of the sidechain nitrogen atoms of the residue in each frame of the trajectory was counted and divided by the total number of frames. (E) Free energy profile of a chloride ion in the WT (blue) and R352Q (red) -CFTR channel pore. The change in free energy relative to site I is plotted versus the chloride ions distance to the alpha carbon of R134 (site III). NBD, nucleotide-binding domain; PM, plasma membrane; R-domain, regulatory domain; TMD, transmembrane domain.

Since the R352 residue lines the CFTR channel pore, the R352Q mutation can be expected to affect the conduction of chloride ions [21]. This was assessed by determining the path of chloride conduction through the channel in WT-CFTR (**Fig 4C, Supplementary material E9**). In the WT system, a chloride ion spontaneously enters the pore intracellular gate [22] and diffuses towards K190 and K370 (site I; **Fig 4C**). It remains in this site for approximately 10 ns and then spontaneously moves up to the vicinity of R352 and R303 (site II). R248, which is positioned in between sites I and II, is involved in coordination of the chloride ion movement between these sites. The chloride ion then moves to the vicinity of R134 and K95 (site III) which coordinate diffusion into the outmost section of CFTR. Chloride conduction through the CFTR channel pore does not follow a straight path from pore entry to site III but rather follows a longer, indirect route. This is in contrast to the straight path that ions follow through voltage-gated potassium and sodium channels, which have tight pores that run perpendicular to the membrane [23].

The role of R352 in chloride conduction was established by comparing the occupancy of the residues along the conduction pathway in the WT- and R352Q-CFTR (**Fig 4D**). In R352Q-CFTR, chloride ions can enter the channel pore but not as readily as in the WT system. They also generally do not progress as far into the pore as for WT-CFTR. Although multiple chloride ions occupied the WT-CFTR channel pore concurrently, no more than one chloride ion at a time was observed in the R352Q-CFTR channel pore. The highest occupancy in the mutant was found for K1041. The extra chloride ions that enter the WT-CFTR channel pore stayed mainly in sites I and II with occupancies of these two sites drastically reduced in R352Q-CFTR compared to WT-CFTR. This reduction in occupancy observed in R352Q-CFTR, is due to neutralisation of the positively charged R352 side chain, which reduces the affinity of chloride ions to site II. Site III was inadequately sampled to draw conclusions (**Fig 4D**). Intriguingly, some chloride ions appeared to penetrate site III in both WT- and R352Q-CFTR, which could explain the mild conductance defect observed for R352Q. Additionally, in simulations of WT-CFTR, it was observed that D993 alternated contact between R303 and R352 (**Supplementary material E8A, B**). However, in R352Q-CFTR – with the replacement of positive R with neutral Q – D993 forms a more stable salt bridge with R303 (**Supplementary material E8C**), which curtails its ability to coordinate a chloride ion. The impact of the lower occupation of sites I and II on chloride conductance was quantified further by comparing the respective free energy profiles obtained from umbrella sampling MD simulations (**Fig 4E**). To generate the initial path for the umbrella sampling simulations, one chloride ion was constrained at site I, and then steered via site II towards R134 in site III, using a moving restraint. The free energy profile calculations followed the protocol described in the **Supplementary material E2**. In WT-CFTR, calculation of the free energy profile yielded an energy trough of −4 kcal/mol at site II relative to site I (**Fig 4E**). This indicates that chloride ions will be attracted to this site. In contrast, in R352Q-CFTR, the free energy profile exhibited an energy barrier of +1 kcal/mol at site II, which will likely hinder chloride conduction. Therefore, in WT-CFTR, a chloride ion in site I will tend to move to site II, whereas such a motion will be suppressed in R352Q-CFTR, resulting in reduced chloride conductance.

## Discussion

This study adds to previously published findings by other CF laboratories by validating the reproducibility of cell models created from CF participants for assessment of CFTR activity via *in vitro* CFTR functional assays [11, 15]. To characterize the rare R352Q-*CFTR* mutation, we first created matched human nasal epithelial cultures (hNECs) and intestinal organoids from participants with *CFTR* mutations that have known functional defects. These acted as references in the functional assays, and since these reference mutations are common to other published platforms [11, 15], this facilitated comparison of data from our functional platform. Our reference hNECs and organoids both showed similar patterns of response to modulators when compared to previous studies, although the magnitude of response is lower [11, 13, 15, 18]. Variabilities in the magnitude of CFTR response in cell models created from different laboratories are evident (**Supplementary material E12**). This could be attributed to several reasons. First, CFTR functional response to modulators in patient cell models were shown to be significantly modified by minor alterations to the culture media [24, 25]. Patient-to-patient variability in CFTR response may also explain this observation [13, 26, 27], which could be further skewed if a relatively small number of participants are studied. The variabilities in data between laboratories highlight the necessity to create reference cell models at every test centre for comparison and characterisation of rare *CFTR* genotypes.

To date, R352Q-CFTR functional characterisation has only been performed in heterologous expression Fischer rat thyroid (FRT) and CFBE41o-cells [28, 29]. These studies demonstrate varying levels of baseline activity ranging from 3-20% of WT-CFTR. However, *in vivo* CFTR function measured using nasal potential difference (NPD) has shown that an individual with the R352Q/G1244E-*CFTR* genotype had CFTR activity just below that of WT individuals [13]. The G1244E mutation is a CFTR gating mutation with severe defect[30], suggesting that R352Q confers high levels of CFTR activity. To better characterize the R352Q mutation, we assessed CFTR function in hNECs and organoids derived from a R352Q/DF508-*CFTR* participant. Of the 100 individuals bearing the R352Q mutation in the CFTR2 database, 99 are heterozygous with DF508 or another *CFTR* mutation genotype [30]. Therefore, our study participant is an appropriate representative of the R352Q-*CFTR* population.

The CFTR activity of a mutation is commonly expressed as a percentage of WT-CFTR activity to measure the severity of defect and responsiveness to modulator [13, 28]. However, we noted limitation in its use because WT-CFTR activity is somewhat underestimated in hNECs, resulting in overestimation of mutant CFTR activity. Baseline CFTR activity in R352Q/DF508 hNECs was 70% of WT-CFTR activity. This was potentiated to well above WT-CFTR activity by VX-770 and G1837. Underestimation of the WT-CFTR activity was also shown by Pranke et al 2017, where VX-770-potentiated CFTR currents for R352Q/G1244E hNECs was approximately 160% of the WT-CFTR activity [13]. In our study, we found that the WT hNECs had significantly higher resting I_sc_ compared to the CF hNECs (WT: 27.5±1.5 μA/cm^2^ vs CF: 1.1±0.7 μA/cm^2^, P<0.0001; data not shown), suggesting that there was endogenous CFTR activity at resting state even without the presence of forskolin. This may have limited further chloride transport when the cells were stimulated with forskolin, underestimating the WT-CFTR activity. The presence of endogenous CFTR activity is also evident in WT intestinal organoids, which were reported to have pre-swollen morphology at resting in the absence of forskolin [11]. Notably, a recent study reported that patient-derived cell models are effective tools to identify modulator-responsive patients but may not accurately predict the magnitude of clinical benefit [31]. It is likely that while the percentage of WT-CFTR activity is useful as a guide of reference for CFTR functional rescue, *in vitro* and *in vivo* CFTR function do not correlate in a linear fashion.

Nonetheless, the high baseline I_sc_ (14.8 μA/cm^2^) and FIS (AUC 196.3) of R352Q indicates a discernible amount of functional CFTR is present at the cell surface. In support of this, VX-809 monotherapy did not further improve CFTR activity in either R352Q/DF508 hNECs or organoids. This is consistent with findings that VX-809 treatment did not further improve VX-770-potentiated CFTR activity in R352Q/G1244E hNECs [13]. Western blot analysis in FRT and CFBE cells showed that 80-98% of R352Q-CFTR is present as mature CFTR localized to the cell surface, with little evidence of a processing or turnover defect [28, 29]. For this reason, because the mechanism of action of VX-809 is correction of a folding defect in CFTR that disrupts its trafficking to the cell surface, it is expected to have little, if any, restorative effect on R352Q-CFTR [32]. Our functional data confirms previous reports that the R352Q mutation is unlikely to cause folding defects and suggests that patients with R352Q-*CFTR* are unlikely to benefit from corrector modulator monotherapy.

Single channel and whole cell patch-clamp studies have demonstrated the positively charged arginine (R352) creates an electropositive potential that promotes anion movement through the channel pore [33]. Additionally, R352 forms a salt bridge with the negatively charged D993 residue (TM9) and this interaction is critical to keep the CFTR channel pore in a stable open-pore state [34]. Our MD simulations performed on the extended structure of human CFTR (PDB: 6MSM) also showed R352Q disrupts the R352-D993 salt bridge in the channel pore, however the gross architecture of the channel pore was unchanged. We showed the R352 residue is not essential for chloride ion entry into the channel pore but is necessary to maintain chloride ion occupancy at a binding site within the channel pore conductance pathway (site II) for normal chloride ion conductivity. This is consistent with previous studies demonstrating R352Q caused frequent transitioning of the channel pore to sub-conductance states (~40-70%) and occasional full conductance channel pore opening [19, 35], causing a mild conductance defect. In the same studies, no apparent reduction in channel pore open probability (Po) was deduced nor reported from the single channel patch-clamp recordings, suggesting no gating defect was present. This is consistent with our MD simulations that predicted the R352Q open conformer can remain stable during chloride ion conduction. This corroborates with the high baseline CFTR activity and significant restoration of CFTR activity by potentiators in R352Q/DF508 hNECs and organoids (close to full WT-CFTR activity levels), in contrast to the G551D/DF508 gating defect reference.

The exact mechanism of action of CFTR potentiators VX-770 and G1837 is not fully understood. There is evidence that VX-770 rescues gating defects by enhancing Po and stabilising the open-pore conformation [36]. VX-770 was initially approved for CFTR gating defects [37]. In some countries, approval has since been extended to conductance defects, based at least in part on *in vitro* functional assays [28, 38]. In principle, potentiators can enhance CFTR activity by increasing the opening time of channels (Po) and/or increasing channel conductance. It is plausible that the significant responsiveness of R352Q/DF508 to potentiators was due to an increase in channel pore opening time. This extra time compensates for the energy barrier created by the R352Q mutation which hinders chloride ion conduction.

Since CF is a multi-organ disease [39], we questioned if the response to modulator therapy in both lung- and intestinal-derived cell models from the same individual would be similar. Overall, our data indicated a positive correlation between CFTR functional response to modulators in matched hNECs and organoids across all *CFTR* mutations assayed. This is consistent with recent findings in several other rare *CFTR* mutations such as G85E, P205S and D614G [40]. In our study, amongst all modulators, VX-809 monotherapy was the only therapy to result in a negative correlation in response between the two cell models in all participants. The hNECs of all participants were responsive to VX-809 monotherapy but not their intestinal organoids. We propose two mechanisms that could result in this discrepancy. First, the VX-809 incubation time of 24 h in organoids is shorter compared to 48 h in hNECs. This could impact on the number of DF508-CFTR which are successfully trafficked from the endoplasmic reticulum to the cell surface since this correction is a time-dependent process [41]. In addition to having a trafficking and gating defect, DF508-CFTR is less stable at the cell surface [42]. Therefore, the shorter VX-809 incubation period in organoids may negatively impact CFTR stabilisation at the cell surface.

Another mechanism could be the different concentration of forskolin used between the two assays. Both the I_sc_ and FIS assays use forskolin to stimulate PKA-dependent CFTR activation [43, 44]. PKA increases channel Po in a dose dependent manner [45]. FIS at 0.128 μM forskolin correlates best with *in vivo* lung function (% predicted FEV_1_) and allows characterisation of responses in organoids without reaching ceiling threshold (saturation) in high residual function *CFTR* mutations [11]. In contrast, 10 μM forskolin has been the default concentration for I_sc_ measurements since early reports of this technique [46, 47]. However, it has recently been shown that 3 μM forskolin is sufficient to saturate the CFTR response in hNECs [48]. The high concentrations of forskolin in the I_sc_ assay are likely to function like a potentiator, extending CFTR channel opening time [45]. This might explain the significant response to VX-809 monotherapy we observed in hNECs but not in matching organoids which were treated with a nearly 100-fold lower concentration of forskolin (0.128 μM).

A strength of our study is the cross-validation of modulator response in matched hNECs and intestinal organoid cultures derived from the same participants. This minimizes the possibility of cell culture-dependent bias. We could not correlate *in vitro* CFTR response to participant *in vivo* clinical response because individuals under 12 years of age with an R352Q-*CFTR* mutation in Australia do not currently have access to CFTR modulator therapy. The combination approach of using *in vitro* experiments and MD simulations to characterize the R352Q-*CFTR* mutation paves a feasible and reliable pathway to personalized therapies for participants with rare *CFTR* mutations. Functional characterisation of CFTR in both lung and intestinal models may improve the specificity and sensitivity of predicting modulator test results and aid in translating patient-derived cell models to become a mainstream companion diagnostic test for people with CF.

## Acknowledgements

We thank the study participants and their families for their contributions. We also thank Sydney Children’s Hospitals (SCH) Randwick respiratory department especially Dr Yvonne Belessis, Leanne Plush, Amanda Thompson and Rhonda Bell in the organisation and collection of participant biospecimens. We thank Nihan Turgutoglu for culture and maintenance of primary cells. SAW is supported by the Australian National Health and Medical Research Council. MA acknowledges support of a top-up scholarship from Cystic Fibrosis Australia. KA is supported by an Australian Government Research Training Program Scholarship. LF is supported by the Rotary Club of Sydney Cove-Sydney Children’s Hospital Foundation and UNSW postgraduate award scholarships. We acknowledge the Hubrecht Institute for the generous provision of L-Wnt3A cell line. Computations were performed on the gadi HPC at the National Computational Infrastructure in Canberra and Artemis at the Sydney Informatics Hub in The University of Sydney.

