## Supplementary material for "Molecular dynamics and theratyping in airway and gut organoids reveal R352Q-CFTR conductance defect"

**Running title**: Personalized medicine in rare cystic fibrosis

Sharon L. Wong[1,2^], Nikhil T. Awatade[1,2^], Miro A. Astore[3^], Katelin M. Allan[1,2], Michael J. Carnell[4], Iveta Slapetova[4], Po-chia Chen[3], Jeffry Setiadi[3], Elvis Pandzic[4], Laura K. Fawcett[1,2,5], John R. Widger[1,2,5], Renee M. Whan[4], Renate Griffith[6,#], Chee Y. Ooi[1,2,7], Serdar Kuyucak[3], Adam Jaffe[1,2,5] and Shafagh A. Waters*[1,2,5]

[1]*School of Women's and Children's Health, Faculty of Medicine, UNSW Sydney, AU.*
[2]*Molecular and Integrative Cystic Fibrosis Research Centre (miCF_RC), UNSW Sydney, AU.*
[3]*School of Physics, University of Sydney, AU.*
[4]*Biomedical Imaging Facility, Mark Wainwright Analytical Centre, UNSW Sydney, AU.*
[5]*Department of Respiratory Medicine, Sydney Children's Hospital, Randwick, AU.*
[6]*School of Chemistry, UNSW Sydney, AU.*
[7]*Department of Gastroenterology, Sydney Children's Hospital, Randwick, AU.*

[^] These authors contributed equally to this work.
[#] Current address: School of Natural Sciences (Chemistry), University of Tasmania

**Supplementary material E1. Characteristics of study participants.**

| Participant ID | CFTR functional defect | *CFTR* genotype | Age (yr) | Sex | Exocrine pancreatic function status | Sweat Chloride (mmol/l) | Nasal epithelial | Intestinal organoids |
|---|---|---|---|---|---|---|---|---|
| CF1 | To be characterised | R352Q/DF508 | 3.1 | M | PS | 79 | Yes | Yes |
| CF2 | Class II – Folding/maturation defect | DF508/DF508 | 4.1 | F | PI | 97 | Yes | Yes |
| CF3 | | DF508/DF508 | 4.1 | F | PI | 92 | Yes | Yes |
| CF4 | | DF508/DF508 | 0.8 | F | PI | Nil | Yes | Yes |
| CF5 | | DF508/DF508 | 1.0 | F | PI | Nil | Yes | Yes |
| CF6 | | DF508/DF508 | 9.0 | M | PI | 90 | Yes | Yes |
| CF7 | Class III – Gating defect | G551D/DF508 | 0.4 | F | PI | 89 | Yes | Yes |
| WT1 | Wild-Type function | WT/WT | 13.3 | M | NA | NA | Yes | No |
| WT2 | | WT/WT | 9.2 | M | NA | NA | Yes | No |
| WT3 | | WT/WT | 3.9 | F | NA | NA | Yes | No |
| WT4 | | WT/WT | 4.0 | M | NA | NA | Yes | No |
| WT5 | | WT/WT | 11.6 | M | NA | NA | Yes | No |
| WT6 | | WT/WT | 0.9 | M | NA | NA | Yes | No |
| WT7 | | WT/WT | 1.2 | M | NA | NA | Yes | No |
| WT8 | | WT/WT | 4.2 | M | NA | NA | Yes | No |
| WT9 | | WT/WT | 6.0 | M | NA | NA | Yes | No |
| WT10 | | WT/WT | 5.7 | M | NA | NA | Yes | No |

PS: pancreatic sufficient; PI: pancreatic insufficient; NA: not applicable; Nil: Sweat chloride was not available.

### Supplementary material E2

#### Materials and methods

#### Study approval and participants biospecimen collection

This study was approved by the Sydney Children's Hospital Ethics Review Board (HREC/16/SCHN/120). Written informed consent was obtained from the legal guardians of all participants. Nasal brushings and/or rectal biopsies were collected during investigative procedures or annual surveillance bronchoscopy from seven CF and ten non-CF participants (**Supplementary material E1**).

#### Nasal epithelial cell culture from nasal brushings

Nasal epithelial cells were collected by brushing the participants inferior nasal turbinates. Using a conditional reprogramming culture (CRC) method, cells were cultured in expansion F-media supplemented with Y-27632 [1]. Briefly, cells were seeded at a density of at least 5,000 cells/cm$^2$ into collagen I-coated (Advanced Biomatrix 5005) flasks, pre-seeded with gamma-irradiated NIH/3T3 feeder cells. Media was changed every second day until cultures reached 80-90% confluence. NIH/3T3 were first dissociated with 1:100 trypsin/EDTA, followed with 1x trypsin/EDTA (Lonza CC-5034) to lift the human Nasal Epithelial Cells (hNECs). These hNECs were now considered passage one and were cryopreserved.

#### Mucociliary differentiation of hNECs at air-liquid interface (ALI)

125,000 passage one hNECs were seeded on the apical side of 6.5mm Transwell® membrane inserts (Sigma CLS3470). Membranes were pre-coated with collagen I. hNECs were expanded for 4-5 days with PneumaCult Ex Plus media (STEMCELL Technologies 05040) applied to both apical and basal compartments. Upon reaching confluence, cultures were switched to ALI culture conditions by complete aspiration of the apical media, exposing the cells to air. Media in the basal compartment was replaced with PneumaCult ALI differentiation media (STEMCELL Technologies 05001) which was replaced every second day for 21-25 days. The presence of beating cilia (ciliogenesis) and mucus production were monitored using light microscopy. From week two post initiation of the ALI culture condition, hNECs were washed with warmed phosphate-buffered saline (PBS) once a week to remove excess mucus.

**Cilia beating frequency measurement**

Cilia beating in differentiated hNECs was imaged and analysed as described previously [1]. Briefly, an ORCA-Flash 4.0 sCMOS camera (Hamamatsu Photonics, Shizuoka Pref., Japan) connected to a Zeiss Axio Observer Z.1 inverted microscope (Carl Zeiss, Jena, Germany) was used to acquire serial images at 334 frames per second (fps) with 3ms exposure time, on a 20x/0.5 objective. All imaging was performed at 37°C, 5% $CO_2$. Image series were analysed using a custom-built script in Matlab (MathWorks, Natick, MA). Image series were filtered to remove the immobile component in each pixel and the Fast Fourier Transform (FFT) algorithm was used to compute the temporal spectrum for each pixel in the image series. The average spectrum per field of view was calculated using the average of all the single pixel spectra. The dominant frequency (highest peak) was then identified using the Matlab function 'findpeaks'.

**Quantification of CFTR-mediated ion transport in differentiated hNECs**

Short circuit current ($I_{SC}$) measurements were performed in the fully differentiated hNECs as described previously [1]. Briefly, Isc recordings were performed under voltage-clamp conditions using VCC MC8 Ussing chambers (Physiologic Instruments, San Diego, CA). Basal-to-apical chloride concentration gradient was created using Ringer solutions with differential composition in the basal and apical chambers. 10 mM HEPES buffered-Ringer solution in basal chamber contained (mM): 145 NaCl, 3.3 $K_2HPO_4$, 10 D-Glucose, 1.2 $MgCl_2$, and 1.2 $CaCl_2$ and apical solution contained (mM): 145 Na-Gluconate, 3.3 $K_2HPO_4$, 10 D-Glucose, 1.2 $MgCl_2$ and 1.2 $CaCl_2$. Ringer solutions were continuously gassed with 95% $O_2$-5% $CO_2$ and maintained at 37˚C. Following a 30 min stabilisation period, hNECs were treated with pharmacological compounds (in order): 100 µM amiloride (apical) to inhibit epithelial sodium channel (ENaC)-mediated $Na^+$ flux, vehicle control 0.01% DMSO or 10 µM VX-770 or 10 µM G1837 (apical) to potentiate cAMP-activated currents, 10 µM forskolin (basal) to induce cAMP activation of CFTR, 30 µM $CFTR_{inh}$-172 (apical) to inhibit CFTR-specific currents and 100 µM ATP (apical) to activate calcium-activated chloride currents. $I_{sc}$ in response to forskolin was considered as baseline activity ($\Delta I_{sc\text{-}Fsk}$) and Isc in response to forskolin and potentiator ($\Delta I_{sc\text{-}Fsk+Pot}$) was used as the measure of modulator response. For assessment of CFTR correction, hNECs were pre-incubated with 3 µM VX-809 for 48h prior to assessment in the Ussing chamber. CFTR modulators are listed in **Supplementary material E10.**

**Whole mount immunofluorescence**

Differentiated hNECs were stained for mucociliary markers including acetylated tubulin, MUC5AC, p63, ZO-1 and E-cadherin as previously described [1]. Briefly, hNECs were fixed in 4% paraformaldehyde or ice-cold methanol-acetone (1:1) for 15 min depending on target protein. Cells were permeabilised with 0.5% Triton-X in PBS on ice for 30 min and blocked using IF buffer (0.1% BSA, 0.2% Triton and 0.05% Tween 20 in PBS) with 10% normal goat serum (Sigma G9023) for 1 h at room temperature before incubation in primary antibodies overnight at 4°C. On the following day, hNECs were washed with IF buffer 3 times, 5 min each and incubated with Alexa Fluor conjugated secondary antibodies for 1 h at room temperature. Cells were mounted with Vectashield hardset antifade mounting medium containing DAPI (H-1500; Vector Laboratories, Burlingame, CA). Images were acquired using

Leica TCS SP8 DLS confocal microscope (Leica Microsystems, Wetzlar, Germany), 63x/1.4 oil immersion objective and images were processed using ImageJ (National Institutes of Health, Bethesda, MD).

**Intestinal organoid culture from rectal biopsies**

Crypts were isolated from four to six rectal biopsies to establish organoid cultures as described previously [2]. Briefly, rectal biopsies were washed with cold PBS. Biopsies were incubated with 10 mM EDTA (Life Technologies 15575-020) in PBS for 120 min on a tube rotator at 4°C. EDTA solution was subsequently removed and crypts were dislodged by vigorous pipetting of biopsies in cold PBS. Isolated crypts were seeded in 70% matrigel (Growth factor reduced, phenol-free; Corning 356231) in 24-well plates at a density of ~10 – 30 crypts in 3x10 µl matrigel droplets per well. The matrigel was polymerised at 37°C for 15 min and then immersed in organoid culture media (**Supplementary material E11**). Media change was performed every second day and organoids were passaged 1:3 after 7 – 10 days of culture. PBS solutions were supplemented with an antibiotic cocktail consisting of vancomycin, gentamicin and fungizone.

**Forskolin-induced swelling assay**

Measurement of fsk-induced swelling (FIS) assay described previously was adapted [3]. Organoids from 7- to 10-day-old culture (passage 3-15) were seeded in flat bottom 96-well culture plates, in 4 µl 70% matrigel droplet per well containing ~25–30 organoids, immersed in 100 µl organoid culture media. The next day, to determine cell viability organoids were incubated with 1.84 µM calcein green (Thermo Fisher Scientific C3100MP) for at least 30 min prior to addition of fsk at 0.02, 0.128, 0.8 or 5 µM concentrations. For CFTR potentiation, 3 µM VX-770 or 3 µM G1837 was added together with fsk. Time-lapse images of organoid swelling were acquired at 10-min intervals for 60 min at 37°C using Zeiss Axio Observer Z.1 inverted microscope (Carl Zeiss, Jena, Germany), on an EC Plan-Neofluar 5x/0.16 M27 dry objective. Where indicated for CFTR correction, organoids were pre-incubated with 3 μM VX-809 for 24 h prior to FIS. Three wells were used per condition and each participant's FIS experiment was repeated 3 to 4 times.

**Quantification of forskolin-induced swelling**

Organoid swelling was quantified using a segmentation strategy implemented using ImageJ/Fiji. Segmentation was performed on brightfield images through classifying pixels based on the pattern of their local neighbourhood with respect to their gradient's magnitude and directionality. The raw image was processed with a gaussian blur (s=1.3) to reduce noise. After the directionality and magnitude of the local gradient was identified, pixels were classified as either 'Background', 'Ridge', 'Valley', 'Rising' or 'Falling' dependent on their neighbouring pixels along the previously calculated local directionality. Clean-up filters were applied that remove noise and small objects, such as ridges that only touched background pixels, and erosions to decrease rising and falling edges to better approximate object boundaries ('Peaks'). Holes were then filled, and a size exclusion was applied that would discriminate debris in the sample preparation from organoids of interest. This segmentation strategy was used to identify area covered by organoid at each time point. The total surface area of organoid at 10-min intervals over 60 min post-fsk stimulation were calculated and normalized against t=0 to render the relative amount of swelling from t=0. The area under the curve, AUC (calculated increase in organoid surface area from t=0 to t=60; baseline=100%) was then calculated using GraphPad Prism software. AUC at 0.8 μm forskolin was considered as the baseline activity and AUC at 0.128 μm forskolin in response corrector and/or potentiator was used as the measure of modulator response.

**Molecular dynamics (MD) system preparation and simulation conditions**

The source of the model was the CFTR structure with PDB ID: 6MSM [4]. We extended this structure to include a section of the regulatory (R) domain (manuscript under preparation). Charges on simulated systems were neutralized with 0.15 M KCl and solvated with TIP3P water [5]. A 1-palmitoyl-2-oleoyl-sn-glycero-3-phosphocholine (POPC) bilayer was generated using the VMD membrane builder plugin in which a model based on the phosphorylated human CFTR channel was embedded [6]. In total 236 lipid molecules and 44503 water molecules were used to build the system. The CHARMM36m forcefield was used for all calculations [7].

**MD simulations of missense mutations**

The 6MSM structure carries the E1371Q-CFTR mutation to prevent hydrolysis of ATP in the nucleotide-binding domain 2 (NBD2), making it easier to capture in the ATP-bound state. This mutation was corrected to match the sequence of human WT-CFTR using the mutator plugin of VMD. The missense mutation R352Q was constructed with the same method. Gromacs v2019.3 was used for all simple MD simulations [8]. Minimisation via a steepest descent algorithm was performed until all forces were below 24 kcal/mol/Å. This was followed by relaxation simulations of all heavy atoms in the system starting with a restraint of 10 kcal/mol/Å$^2$ and then halving this restraint every 200 ps in 15 iterations. Relaxation and production were run with 1 and 2 fs time steps respectively. Relaxation was followed by 5 ns of equilibration. The z axis of the system was normal to the membrane, after relaxation the area in the x-y plane was fixed for the production run. During relaxation, a Berendsen thermostat and barostat were applied, and for production we switched to a Nosé-Hoover and Parrinello-Rahman thermostat and barostat, respectively [9-11]. Production runs were extended up to 2 μs at 310K with three replicates for all simple MD simulations. All covalent bonds involving hydrogen were constrained using the LINCS algorithm [12]. The last microsecond of the longest simulations for each system were selected for further analysis. All root-mean-square deviations (RMSDs) were calculated using the positions of alpha carbons with reference to the 6MSM experimental structure [4]. Analysis scripts were written in python using the MDAnalysis library and are available online at https://github.com/miro-astore/mdanalysis_scripts [13, 14].

#### Free energy calculations

NAMD v2.13 was used for the umbrella sampling MD simulations to determine the free energy surface for chloride ion conductance through a section of the channel pore [15]. The proposed chloride ion pathway (Fig 4C) was assessed with umbrella sampling. Starting from site I, at a distance of 33.5 Å from the alpha carbon of R134, the chloride ion was pulled further into the channel pore with a spring force of 10 kcal/mol/Å$^2$. The chloride ion was pulled up to 19 Å from the alpha carbon of R134 and stratified at 0.5 Å intervals. This reference point was chosen because R134 is the highest positive amino acid before the constriction in the 6MSM structure and the furthest the chloride ion was observed to travel in the unbiased MD simulations. Each umbrella window was run for 70 ns for both the WT and R352Q systems. The free-energy profile (**Fig 4E**) was obtained using the weighted histogram analysis method [16]. The profile is compiled as the average from five 7 ns blocks of samples with uncertainties given by the standard error of the mean (SEM) of the five samples. A temperature of 300K was used for the umbrella sampling simulations.

#### Statistical analysis

Data for **Fig 1B** and **Supplementary material E7** are presented as dot plots with mean ± standard error of the mean (SEM). Data in **Fig 1D, 2C** and **Supplementary material E3** are presented as violin plots with mean and all data points presented. The mean ± SEM for **Fig 1D** and **2C** are presented in tables in **Supplementary material E5** and **S6**. Data for **Fig 2A** are presented as line graphs with mean ± standard deviation (SD). One-way analysis of variance (ANOVA) was used to determine statistical differences. Pearson correlation coefficient was used to determine association between modulator-stimulated CFTR response (corrected for baseline CFTR activity) in hNECs $I_{SC}$ measurements compared to FIS response in intestinal organoids. Statistical analysis was performed with GraphPad Prism software v9.0.1. A *P*-value of less than 0.05 was considered to be statistically significant.

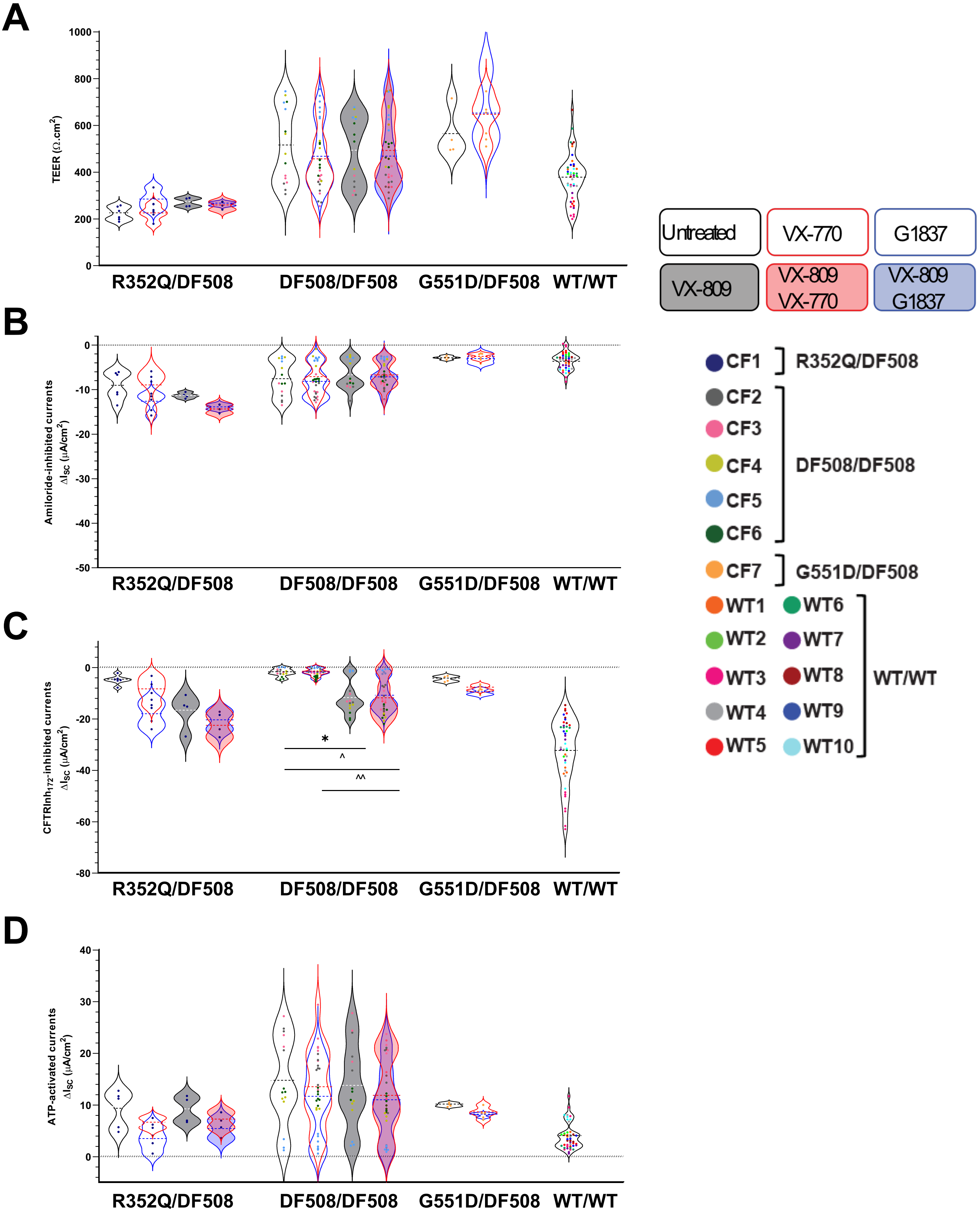


**Supplementary material E3. Electrophysiological parameters of differentiated hNECs from one R352Q/DF508, five DF508/DF508, one G551D/DF508 participants with CF and ten WT-CFTR control participants. (A)** Trans-epithelial electrical resistance (TEER). **(B)** Amiloride-inhibited epithelial sodium channel (ENaC) currents. **(C)** $CFTR_{inh}$-172-inhibited currents. **(D)** ATP-stimulated calcium-activated chloride channel (CaCC) currents. Each participant is coded with a different colour. Each dot represents an individual replicate. Data represented as violin plots to show distribution of data. One-way analysis of variance (ANOVA) was used to determine statistical differences. *P < 0.05. ^P < 0.05 and ^^P < 0.01 for VX-770 only.

**Supplementary material E4. Functional beating cilia in differentiated hNECs from a DF508/DF508 participant (CF3).** Online video

### Supplementary material E5. Short circuit current (Isc) measurements in response to modulators for hNECs from all *CFTR* genotypes

| *CFTR* genotype | **WT/WT n=10** |
|---|---|
| Treatment | **Untreated (DMSO-baseline)** |
| Mean ΔIsc-fsk | 21.2 |
| SEM | 1.2 |

| *CFTR* genotype | **DF508/DF508 n=5** | | | | | | **G551D/DF508 n=1** | | | **R352Q/DF508 n=1** | | | | | |
|---|---|---|---|---|---|---|---|---|---|---|---|---|---|---|---|
| Treatment | **Untreated (DMSO-baseline)** | **VX-770** | **G1837** | **VX-809** | **VX-809 + VX-770** | **VX-809 + G1837** | **Untreated (DMSO-baseline)** | **VX-770** | **G1837** | **Untreated (DMSO-baseline)** | **VX-770** | **G1837** | **VX-809** | **VX-809 + VX-770** | **VX-809 + G1837** |
| Mean ΔIsc-fsk | 3.4 | 3.9 | 4.1 | 12.4 | 13.8 | 14.3 | 4.2 | 8.4 | 12.4 | 14.8 | 30.0 | 38.3 | 23.9 | 32.4 | 40.6 |
| SEM | 0.5 | 0.4 | 0.5 | 1.5 | 1.5 | 1.8 | 0.8 | 0.5 | 1.1 | 1.4 | 2.3 | 3.4 | 3.2 | 6.3 | 2.3 |
| Absolute difference to Untreated | - | 0.5 | 0.7 | 9.0 | 10.4 | 10.9 | - | 4.2 | 8.2 | - | 15.2 | 23.5 | 9.1 | 17.6 | 25.8 |
| Fold change to Untreated | - | 1.2 | 1.2 | 3.7 | 4.1 | 4.2 | - | 2.0 | 3.0 | - | 2.0 | 2.6 | 1.6 | 2.2 | 2.7 |
| Absolute difference to VX-770 | - | - | - | - | 9.9 | - | - | - | - | - | - | - | - | 2.4 | - |
| Absolute difference to G1837 | - | - | - | - | - | 10.1 | - | - | - | - | - | - | - | - | 2.3 |
| Contribution of one DF508 allele to ΔIsc-fsk | 3.4/2=1.7 | 0.5/2=0.25 | 0.7/2=0.35 | 9.0/2= 4.5 | 10.4/2=5.2 | 10.9/2=5.5 | - | - | - | - | - | - | - | - | - |

SEM = standard error of the mean

### Supplementary material E6. Forskolin-induced swelling (FIS) in response to modulators for organoids from all *CFTR* genotypes

| | DF508/DF508 n=5 | | | | | | G551D/DF508 n=1 | | | | | | R352Q/DF508 n=1 | | | | | |
|---|---|---|---|---|---|---|---|---|---|---|---|---|---|---|---|---|---|---|
| | **Untreated (DMSO-baseline)** | **VX-770** | **G1837** | **VX-809** | **VX-809 + VX-770** | **VX-809 + G1837** | **Untreated (DMSO - baseline)** | **VX-770** | **G1837** | **VX-809** | **VX-809 + VX-770** | **VX-809 + G1837** | **Untreated (DMSO - baseline)** | **VX-770** | **G1837** | **VX-809** | **VX-809 + VX-770** | **VX-809 + G1837** |
| Mean AUC | 27.3 | -2.3 | 29.4 | 12.3 | 187.0 | 175.0 | 67.1 | 106.0 | 121.3 | 97.5 | 95.6 | 110.0 | -47.3 | 187.3 | 863.3 | -29.6 | 542.5 | 1323.0 |
| SEM | 18.7 | 17.8 | 14.6 | 14.5 | 29.2 | 22.2 | 16.9 | 22.4 | 32.4 | 25.5 | 23.9 | 30.5 | 10.6 | 15.5 | 49.2 | 10.0 | 57.9 | 90.2 |
| Absolute difference to Untreated | - | -29.6 | 2.1 | -15.0 | 159.7 | 147.7 | - | 38.9 | 54.2 | 30.4 | 28.5 | 42.9 | - | 234.6 | 910.6 | 17.7 | 589.8 | 1370.3 |
| Absolute difference to VX-770 | - | - | - | - | 189.3 | - | - | - | - | - | -10.4 | - | - | - | - | - | 355.2 | - |
| Absolute difference to G1837 | - | - | - | - | - | 145.6 | - | - | - | - | - | -11.3 | - | - | - | - | - | 459.7 |
| Contribution of one DF508 allele to AUC | 27.3/2=13.7 | -29.6/2=-14.8 | 2.1/2=1.1 | -15.0/2=-7.5 | 159.7/2=79.9 | 147.7/2=73.9 | - | - | - | - | - | - | - | - | - | - | - | - |

SEM = standard error of the mean

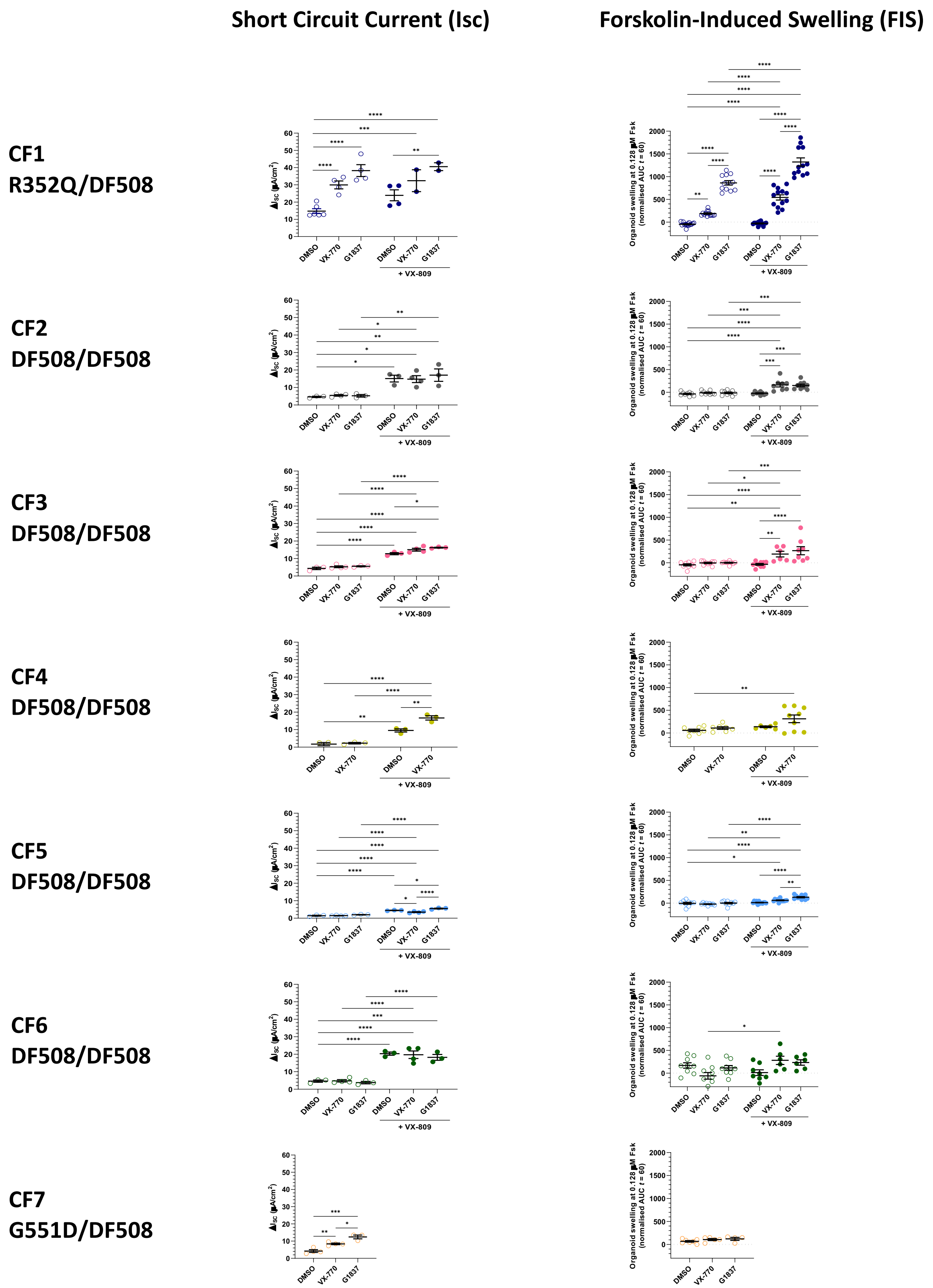


**Supplementary material E7. Paired total stimulated currents, $\Delta I_{sc}$ and forskolin-induced swelling, AUC at 0.128 µM fsk of individual participant CF1 to CF7.** Data are means ± SEM from at least three treatment replicates (each dot represents an individual replicate). One-way analysis of variance (ANOVA) was used to determine statistical differences. *P < 0.05, **P < 0.01, ***P < 0.001 and ****P < 0.0001.

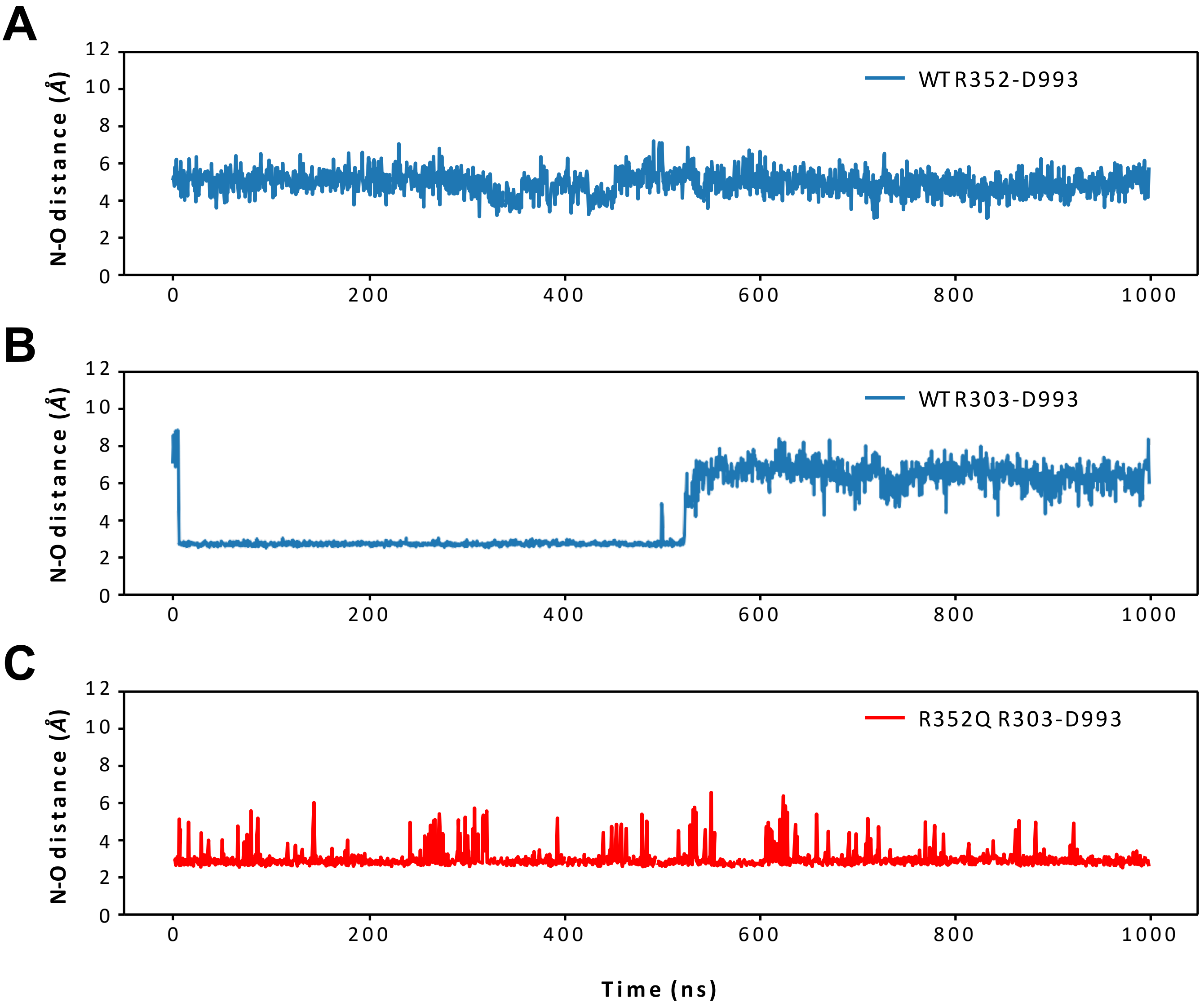


**Supplementary material E8. The stability of key salt bridges within the CFTR channel.** N-O distance refers to the distance between the charged head groups of the positive arginine (R303, R352) residue and the negative aspartate (D993) residue. **(A)** It is expected that a stable interaction between R352-D993 is an important feature of the fully open channel. Our data indicates that R352 remains in close proximity to D993 in the Wild Type channel. **(B)** R303 makes a stable salt bridge with D993 for the first 550 ns of the simulation. The presence of R352 allows D993 to either interact with R352 or R303. When D993 is in contact with R352, D993 is neutralised, leaving R303 free to coordinate a nearby chloride ion. **(C)** In the R352Q mutant, the R303-D993 salt bridge remains throughout the simulation.

**Supplementary material E9. Unbiased simulation of chloride ion conductance in WT- and R352Q-CFTR**. Online video

**Supplementary material E10. Modulator compounds used for hNECs Isc measurement and organoid FIS assay.**

| Modulator | Supplier | Stock concentration | Final concentration (FIS) | Final concentration (Isc) | Solvent | Storage |
|---|---|---|---|---|---|---|
| VX-809 | Selleckchem S1565 | 10 mM | 3 µM | 3 µM | DMSO (Sigma D2650) | -80ºC |
| VX-770 | Selleckchem S1144 | 10 mM | 3 µM | 10 µM | DMSO (Sigma D2650) | -80ºC |
| GLPG1837 | Selleckchem S8698 | 10 mM | 3 µM | 10 µM | DMSO (Sigma D2650) | -80ºC |

The concentration and incubation time of CFTR modulators are the same as previous published studies [17, 18]. GLPG1837 was used at the same concentration as VX-770 for consistency.

**Supplementary material E11. Components of intestinal organoid culture media.**

| **Components** | **Concentrations** | **Supplier** |
|---|---|---|
| Advanced DMEM/F-12 media | - | Life Technologies 12634-010 |
| Penicillin-Streptomycin | 1x | Sigma P4333 |
| HEPES | 10 mM | Sigma H0887 |
| L-glutamine | 2 mM | Sigma G8541 |
| B-27 Supplement | 1x | Life Technologies 17504-044 |
| N-acetylcysteine | 1.25 mM | Sigma A9165 |
| hEGF | 50 ng/ml | Sigma E9644 |
| Wnt3a-conditioned media (WCM) | 50% | In-house [3] |
| hRspo-1 | 300 ng/ml | Peprotech 120-38 |
| hNoggin | 100 ng/ml | Peptrotech 120-10C |
| Nicotinamide | 10 mM | Sigma N0636 |
| A83-01 | 500 nM | Tocris Bioscience 2939 |
| SB202190 | 10 µM | Sigma S7067 |
| Gentamicin | 50 µg/ml | Sigma G1397 |
| Vancomycin | 50 µg/ml | Sigma V2002 |
| Fungizone | 250 ng/ml | Life Technologies 15290-018 |
| Primocin | 100 µg/ml | InvivoGen ant-pm-2 |

Abbreviations: hEGF, human epidermal growth factor.

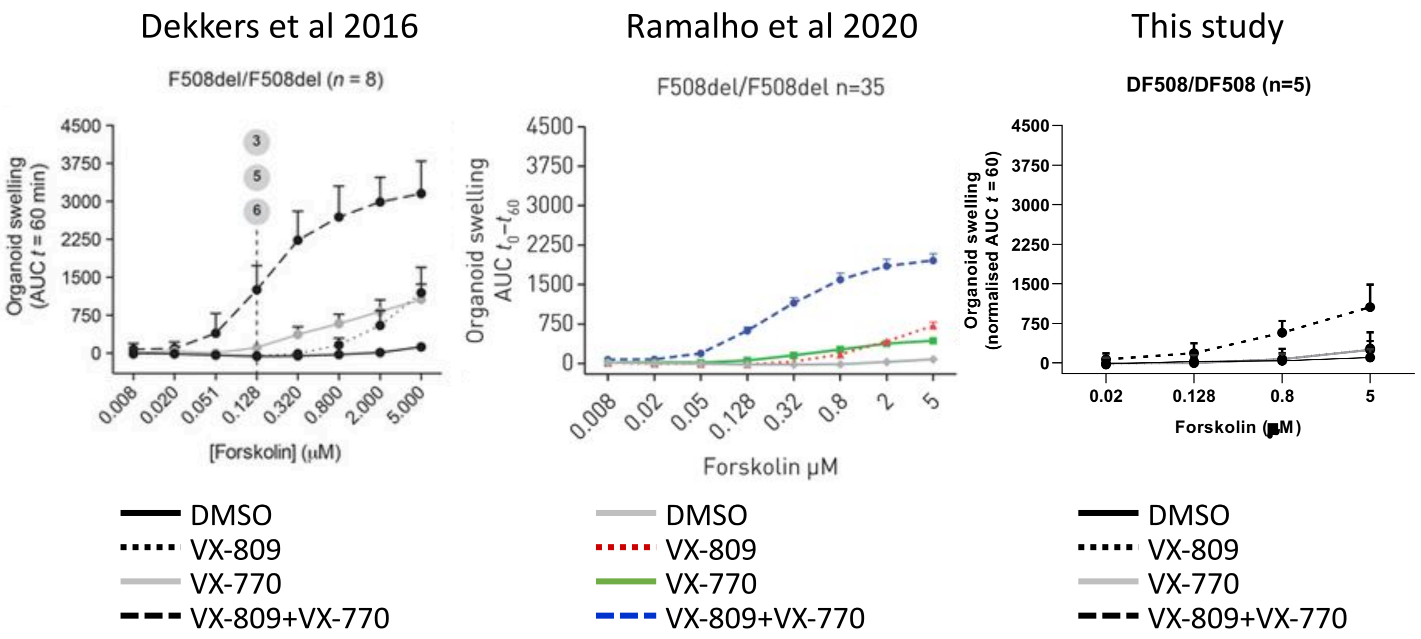


**Supplementary material E12. Comparison of forskolin-induced assay data in DF508/DF508 intestinal organoids from three different laboratories.** Graphs on the left and middle were reproduced from [18] and [19] respectively with copyright permission.
